# Chromosome-scale assembly of the bread wheat genome, *Triticum aestivum*, reveals over 5700 new genes

**DOI:** 10.1101/2020.04.06.028746

**Authors:** Michael Alonge, Alaina Shumate, Daniela Puiu, Aleksey Zimin, Steven L. Salzberg

## Abstract

Bread wheat (*Triticum aestivum)* is a major food crop and an important plant system for agricultural genetics research. However, due to the complexity and size of its allohexaploid genome, genomic resources are limited compared to other major crops. The IWGSC recently published a reference genome and associated annotation (IWGSC v1.0, Chinese Spring) that has been widely adopted and utilized by the wheat community. Although this reference assembly represents all 3 wheat subgenomes at chromosome scale, it was derived from short reads, and thus is missing a substantial portion of the expected 16 gigabases of genomic sequence. We earlier published an independent wheat assembly (Triticum 3.1, Chinese Spring) that came much closer in length to the expected genome size, although it was only a contig-level assembly lacking gene annotations. Here, we describe a reference-guided effort to scaffold those contigs into chromosome-length pseudomolecules, add in any missing sequence that was unique to the IWGSC 1.0 assembly, and annotate the resulting pseudomolecules with genes. Our updated assembly, Triticum 4.0, contains 15.07 gigabases of non-gap sequence anchored to chromosomes, which is 1.2 gigabases more than the previous reference assembly. It includes 108,639 genes unambiguously localized to chromosomes, including over 2000 genes that were previously unplaced. We also discovered more than 5700 new genes, all of them duplications in the Chinese Spring genome that are missing from the IWGSC assembly and annotation. The Triticum 4.0 assembly and annotations are freely available at www.ncbi.nlm.nih.gov/bioproject/PRJNA392179.

## INTRODUCTION

Bread wheat (*Triticum aestivum*) is a crop of significant worldwide nutritional, cultural and economic importance. As with most other major crops, there is a strong interest in applying advanced breeding and genomics technologies towards crop improvement. Key to these efforts are high-quality reference genome assemblies and associated gene annotations which are the foundations of genomics research. However, the bread wheat genome has some notable features that make it especially technically challenging to assemble. One such feature is allohexaploidy (2n=6×=42, AABBDD), a result of wheat’s dynamic domestication history^1^. Specifically, this polyploidy results from the hybridization of domesticated emmer (*Triticum turgidum*, AABB) with *Aegilops tauschii* (DD). Domesticated emmer, also an ancestor of durum wheat, is itself an allotetraploid resulting from interspecific hybridization between *Triticum* and *Aegilops* species.

The resulting bread wheat genome is immense, with flow cytometry studies estimating the genome size to be ∼16 Gbp^2^. As with most other large plant genomes, repeats, especially retrotransposons, make up the majority of the genome, which is estimated to be ∼85% repetitive^3^. Taken together, these genomic characteristics make for an especially difficult genome to assemble, even given the recent improvements in long-read sequencing and algorithmic advancements in genome assembly technology. Nonetheless, early efforts were made to establish *de novo* reference genome assemblies. In 2014, the International Wheat Genome Sequencing Consortium (IWGSC) used flow cytometry-based sorting to sequence and assemble individual chromosome arms, thus removing the repetitiveness introduced by homeologous chromosomes^4^. In spite of this approach, this short-read based assembly was highly fragmented, and only reconstructed ∼10.2 Gbp of the genome. Subsequent short-read assemblies using alternate strategies were also developed by the community, though each also struggled to achieve contiguity and completeness^5,6^.

In 2017, we released the first-ever long-read-based assembly for bread wheat (Triticum 3.1), representing the Chinese Spring variety^7^. With an N50 contig size of 232.7 kb, Triticum 3.1 was far more contiguous than any previous assembly of bread wheat, and with a total assembly size of 15.34 Gbp, it reconstructed the highest percentage of the expected wheat genome size of any assembly. Though this assembly provided a more complete representation of the Chinese Spring genome, its contigs were not mapped onto chromosomes, and notably, it did not include gene annotation.

In 2018, the IWGSC published a chromosome-scale reference assembly and associated annotations for bread wheat (v1.0, Chinese Spring), providing the best-annotated reference genome yet^3^. Because that assembly was entirely derived from short reads, it was less complete and more fragmented then Triticum 3.1, having a total size of 14.5 Gb and an N50 contig size of 51.8 Kb. However, a collection of long-range scaffolding data, including physical (BACs, Hi-C), optical (Bionano), and genetic maps, enabled most of the assembled scaffolds to be mapped onto wheat’s 21 chromosomes. These pseudomolecules served as a foundation for comprehensive *de novo* gene and repeat annotation, facilitating investigations into the genomic elements that drove the evolution of genome size, structure, and function in wheat.

Here, we used the IWGSC v1.0 assembly (Genbank accession GCA_900519105.1 to inform the scaffolding and annotation of the more complete Triticum 3.1 assembly. The new assembly, Triticum 4.0, contains 1.1 Gbp of additional non-gapped sequence compared to IWGSC v1.0, while localizing 97.9% of sequence to chromosomes. Comparative analysis revealed that Triticum 4.0 more accurately represents the Chinese Spring repeat landscape, which is heavily collapsed in IWGSC v1.0. Our more-complete assembly allowed us to anchor ∼2,000 genes that were previously annotated on unlocalized contigs in IWGSC v1.0. We also found 5,799 additional gene copies in Triticum 4.0, showing extensive collapsing of gene duplicates in IWGSC v1.0 assembly. We highlighted one such example mis-assembly to demonstrate that potentially functionally relevant genes are represented with improved accuracy in Triticum 4.0. The Triticum 4.0 assembly and annotations are available without restriction at www.ncbi.nlm.nih.gov/bioproject/PRJNA392179.

## RESULTS

### Scaffolding the Triticum 3.1 genome assembly

Our goal was to utilize both our previously published Triticum 3.1 contigs (T3) and the IWGSC v1.0 assembly (IW) to establish an improved chromosome-scale genome assembly for the Chinese Spring variety of bread wheat. **Figure 1** depicts the pipeline used to derive our final Triticum 4.0 (T4) assembly. As a foundation, we started with the T3 contigs because they were highly contiguous (N50 = 232.7Kb) and contained a total of 1.1Gbp more non-gap sequence compared to the IW assembly. However, we wanted to ensure that our final assembly did not exclude any sequence missing from T3 but present in IW. To incorporate any such “missing” IW contigs, we first derived a set of contigs from the IW assembly by breaking pseudomolecules at gaps. By aligning these IW contigs to the T3 assembly, we identified 4,702 IW contigs (89,866,936 bp) with sequence missing from the T3 assembly. These sequences along with the T3 contigs comprised our initial contig set.

**Figure 1.**
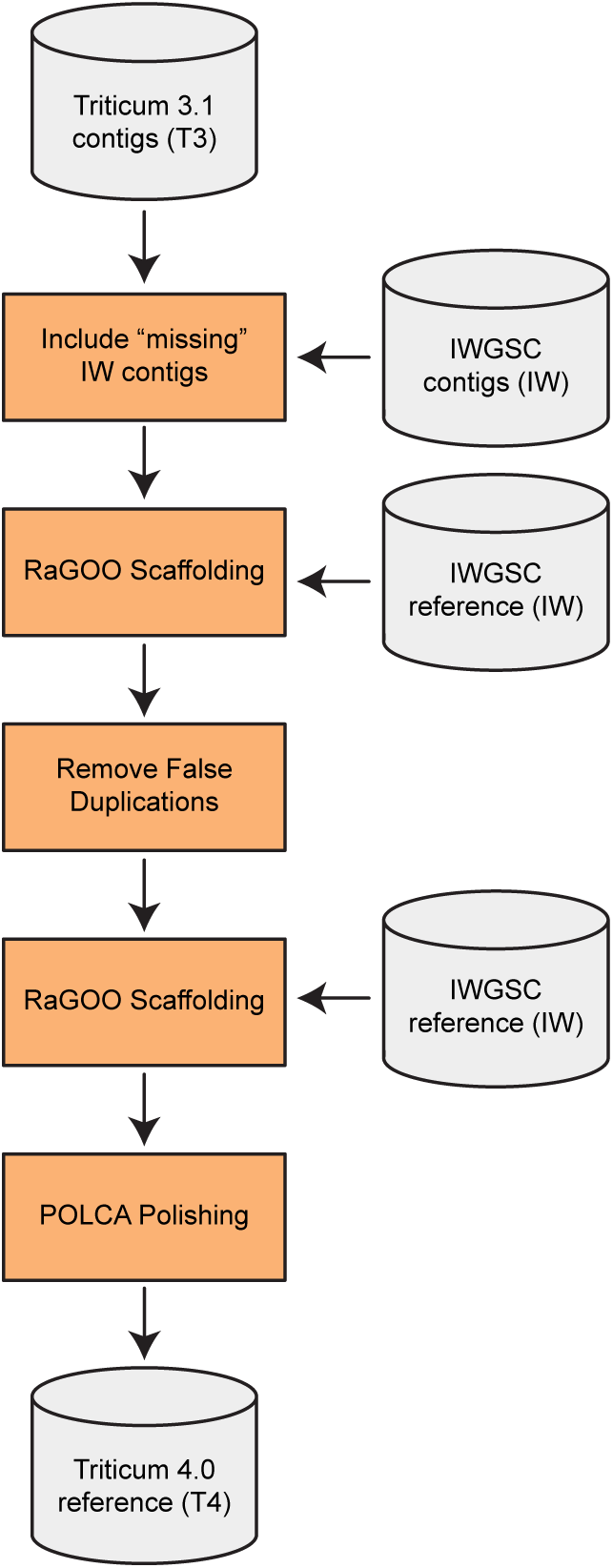
The Triticum 4.0 assembly scaffolding pipeline. A diagram depicting the Triticum 4.0 assembly scaffolding pipeline. Grey cylinders represent input or output genome assemblies, while orange boxes show the steps of the scaffolding process.

In order to create chromosome-length scaffolds (pseudomolecules), we used RaGOO^8^ to perform reference-guided scaffolding on our input contigs. This scenario presents a near-ideal context for reference-guided scaffolding, because the contigs and the reference assembly represent the same inbred genotype, and thus no genomic structural differences are expected. Although RaGOO normally utilizes Minimap2^9^ alignments between contigs and a reference assembly, we used NUCmer^10,11^ instead, as it offered the necessary flexibility to align these large and repetitive genomes. Specifically, NUCmer provided the specificity needed to unambiguously align contigs to a highly repetitive allohexaploid reference genome (see **Methods**). Even with these high stringency alignments, a majority of sequence (97.67% of bp) was ordered and oriented into pseudomolecules.

We next sought to remove any false duplications potentially created during the process of incorporating 4,702 IW sequences. We aligned these IW contigs to the RaGOO scaffolds and removed 357 IW contigs from the initial set of 4,702 that aligned to more than one place in the assembly and therefore were no longer deemed “missing” from T3. This produced our final set of contigs, which included the T3 contigs plus 4,345 (84,909,842 bp) contigs from IW that contained sequence missing from T3. The final contigs had an N50 length of 230,687 bp (essentially the same as the T3 assembly) and a sum of 15,429,603,425 bp. We then repeated the RaGOO scaffolding step, and the resulting scaffolds were polished with POLCA^12^ using the original Illumina reads, yielding the final T4 chromosome-scale assembly.

Despite the highly repetitive nature of the genome, RaGOO confidence scores indicate that T4 scaffolding was accurate and unambiguous (**Figure S1**). This suggests that our high-specificity NUCmer parameters mitigated erroneous contig ordering and orientation resulting from repetitive alignments. Accordingly, dotplots confirm that, as expected, there are no large-scale structural rearrangements between T4 and IW pseudomolecules (**Figure S2**). While borrowing its chromosomal structure from IW, T4 demonstrates superior sequence completeness. 97.9% of T4 sequence (15.09 Gbp) was placed onto 21 chromosomes yielding pseudomolecules that had 1.2 Gbp more localized non-gapped sequence than the IW reference (**Table 1**). This extra sequence was distributed evenly across the genome, with each T4 pseudomolecule containing more sequence (average of 48.8 +/- 8.4 Mbp) than its IW counterpart while having substantially fewer gaps (**Figure 2**).

**Table 1.** Non-gapped sequence length of the Triticum 4.0, IWGSC v1.0, and Triticum 3.1 assemblies.

| **Assembly** | **Triticum 4.0** | **IWGSC v1.0** | **Triticum 3.1** |
| --- | --- | --- | --- |
| All sequence (bp) | **15,397,713,314** | 14,271,578,887 | 15,344,693,583 |
| Anchored sequence (bp) | **15,070,919,678** | 13,840,498,961 | N/A |

**Figure 2.**
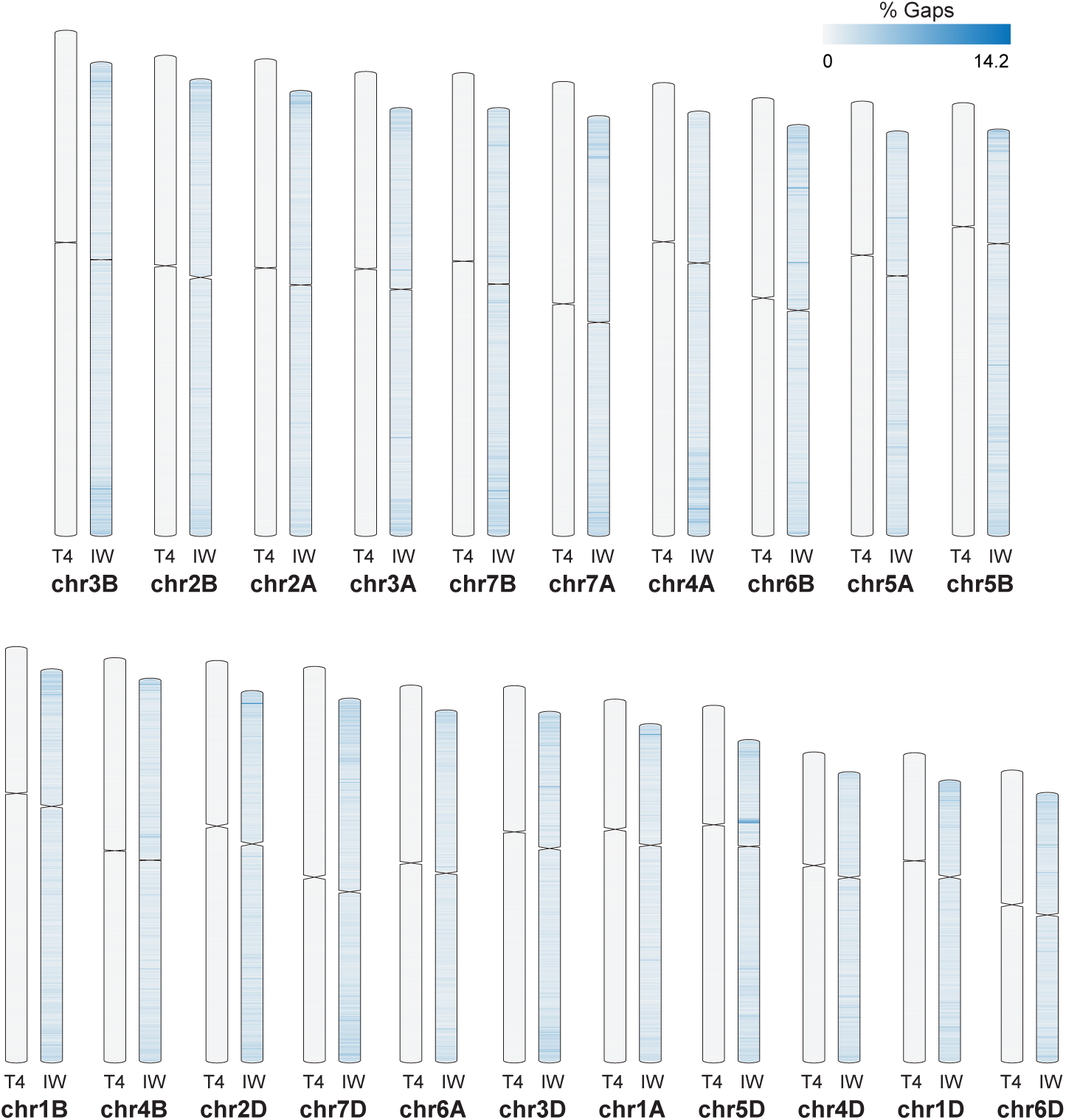
A comparison of T4 and IW assembly completeness. An ideogram showing the distribution of gap sequence in the Triticum 4.0 (T4) and IWGSC v1.0 (IW) assemblies. The heatmap color intensity corresponds to the percentage of gap sequence in non-overlapping 1 Mbp windows along each chromosome. Each T4 chromosome across all 3 subgenomes has more sequence and fewer gaps than its IW counterpart.

Since IW was derived from short-reads, it is conceivable that some genomic repeats were collapsed during assembly^13^. Therefore, we hypothesized that T4, a long-read-based assembly, more accurately represents the repeat landscape of the Chinese Spring genome. As support for this hypothesis, we observe that 101-mers shared by T4 and IW were present at higher copies in T4 (**Figure 3**). This observation holds for a wide range of 101-mer copy numbers, suggesting that T4 more accurately represents both lower-order (duplications) and higher-order (transposable elements) repeats. To investigate a specific instance of repeat collapse in IW, we compared centromere sequence content in the two assemblies. As was done in the original IW publication, we used publicly available CENH3 ChIP-seq data to define centromere positions in T4 (see **Methods**) (**Table S1**)^3^. This analysis indicated ChIP-seq peaks corresponding to centromeres for each of the 21 chromosomes (**Figure S3**). Notably, T4 had a total of 38.8 Mbp more centromeric sequence than IW, highlighting that the long-read-based T4 assembly collapsed less centromeric sequence than IW.

**Figure 3.**
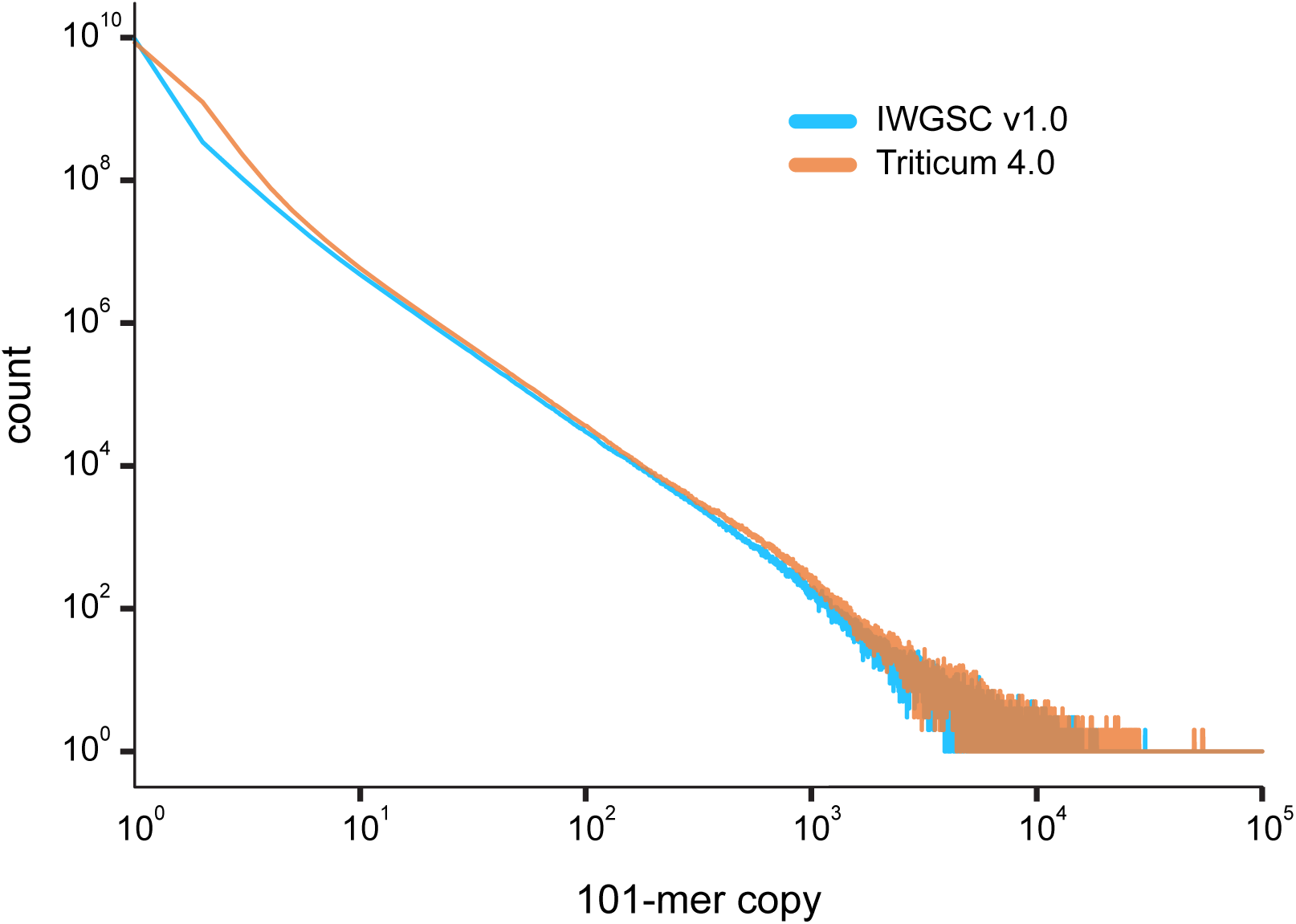
Shared assembly k-mer count distribution. Histogram of 101-mer copy number in the Triticum 4.0 and IWGSC v1.0 assemblies. Only 101-mers shared by both assemblies are considered. While IW has more single-copy 101-mers, T4 represents more 101-mers at higher copy numbers.

### Annotating the Triticum 4.0 genome assembly

We mapped the IW v1.1 high-confidence annotation onto T4 using an annotation lift-over tool we developed called Liftoff (see **Methods**) (https://github.com/agshumate/Liftoff). Given a genome annotation, Liftoff aligns all genes, chromosome by chromosome, to a different genome of the same species using BLAST^14^. For all genes that fail to map to the same chromosome, Liftoff attempts to map them across chromosomes. The best mapping for each gene is chosen according to sequence identity and concordance with the exon/intron structure of the original gene model. Out of 130,745 transcripts from 105,200 gene loci annotated on primary chromosomes in IW, we successfully mapped 124,711 transcripts from 100,839 gene loci. We define a transcript as successfully mapped if the mRNA sequence in T4 is at least 50% as long as the mRNA sequence in IW. However, the vast majority of transcripts greatly exceed this threshold, with 92% percent of transcripts having an alignment coverage of 98% or greater (**Figure S4A**). Sequence identity is similarly high with 92% of transcripts aligning at an identity of 95% or greater (**Figure S4B**). Of the transcripts that failed to map, 4,888 had a partial mapping with an alignment coverage < 50%, and the remaining transcripts failed to map entirely.

The IW annotation also contains 2,691 genes annotated on unplaced contigs (called chrUn in IW v1.1). Using Liftoff, we were able to map 2,001 of these genes onto a primary chromosome in T4, the majority of which (68%) aligned with 100% coverage and identity (**Table S2**). To control for differences in annotation pipelines between IW and T4, we used Liftoff to map chrUn genes onto the primary IW chromosomes to look for additional, unannotated gene copies. Of the 2,001 chrUn genes mapped to T4 pseudomolecules, 78 of these were also mapped to primary IW chromosomes. This suggests that at least 1,923 genes were placed due to improved assembly completeness rather than differences in annotation methods.

After mapping the IW v1.1 annotation onto T4, we used Liftoff to look for additional gene copies in T4. We required 100% alignment coverage and 100% sequence identity in exons and splice sites to map a gene copy. We found 5,799 additional gene copies in T4 that are not annotated in IW v1.1 (**Table S3**). 4,158 genes have 1 extra copy and 567 genes have 2 or more additional copies, with a maximum of 84 additional copies (**Figure 4A**). IW collapsed most genes copies on the same chromosome rather than across homeologous chromosomes, with 4,062 of the 5,799 additional gene copies occurring on the same chromosome and 97 copies occurring on the same chromosome of a different subgenome (**Figure 4B**). 915 gene copies were placed on different chromosomes. The remaining 725 are extra copies of chrUn genes placed on chromosomes. As was done for unplaced genes, we also looked for additional IW gene copies present elsewhere in IW. Of our 5,799 additional gene copies, 159 were also present in IW suggesting that at least 5,640 of T4 copies are strictly the result of improved assembly completeness.

**Figure 4.**
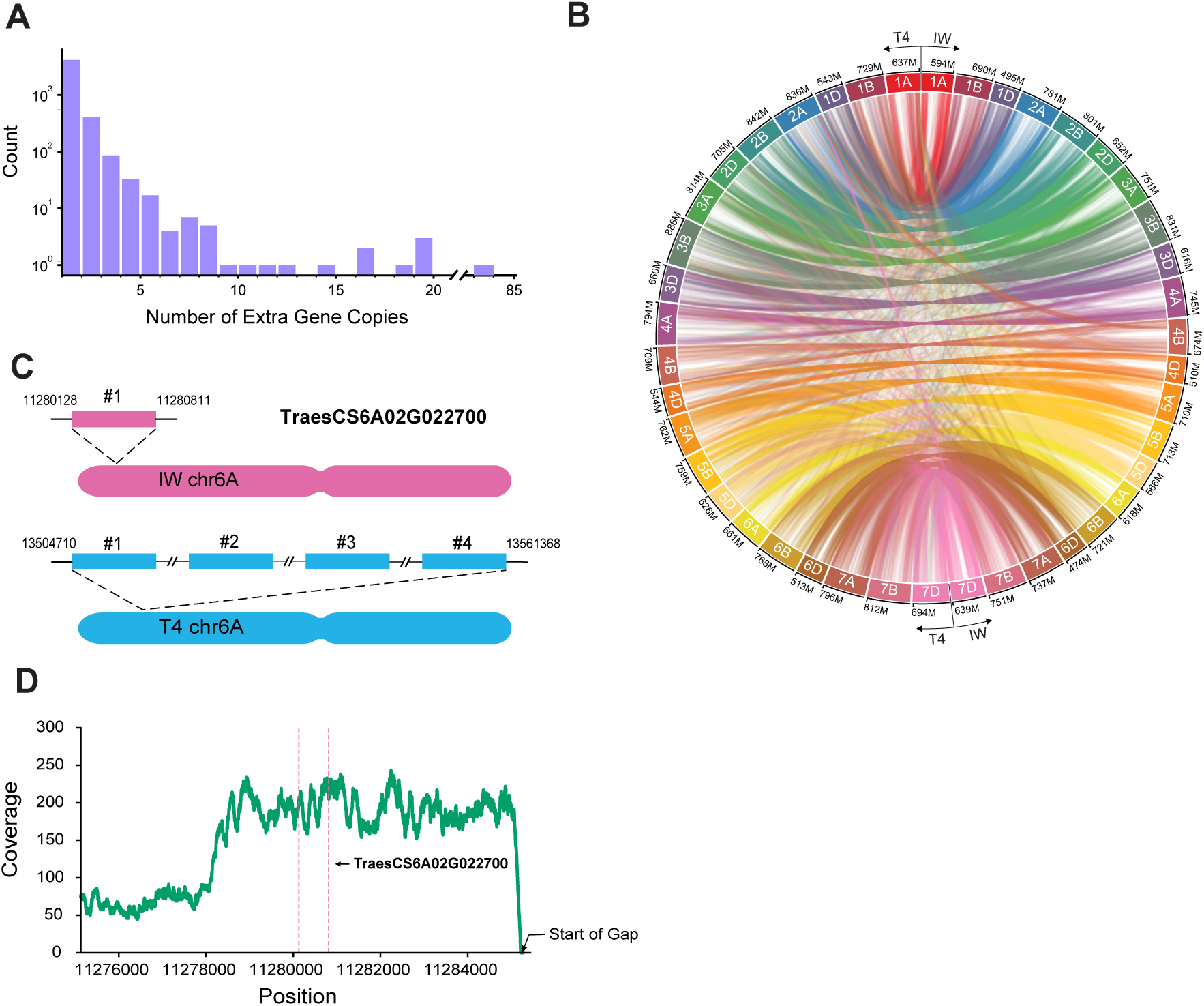
Triticum 4.0 resolves previously collapsed genic repeats. **(A)** Histogram depicting the distribution of the number of additional gene copies found in Triticum 4.0 (T4). **(B)** Circos plot showing the locations of all additional gene copies (http://omgenomics.com/circa/). Lines are drawn from the location on the gene in IWGSC v1.0 (IW) on the right half of the diagram to the location of each copy in T4 on the left half. **(C)** Diagram depicting the MADS-box transcription factor gene, *TraesCS6A02G022700*, present in 3 additional tandem copies in T4 as relative to IW. Ideograms are not drawn to scale. **(D)** Plot of the short-read coverage in IW starting 5kb upstream of *TraesCS6A02G02270* and extending to the first gap downstream of the gene. The pink dashed lines show the location of the gene.

We looked for specific examples of potentially functionally relevant gene duplications previously collapsed in IW. One such example was a MADS-box transcription factor gene, *TraesCS6A02G022700*, which we found with three additional tandem copies on chr6A in T4 relative to IW (**Figure 4C**). MADS-box transcription factors are known to be regulators of plant development, influencing traits such as flowering time and floral organ development^15,16^. Furthermore, it has been demonstrated that MADS-box genes can quantitatively impact gene expression and domestication phenotypes in a dosage dependent manner^17^. To provide further evidence that this gene is part of a collapsed repeat in IW, we aligned Chinese Spring Illumina reads to IW and calculated the coverage across the gene +/- 50kb of flanking sequence. We observed a spike in coverage indicating a collapsed repeat in IW containing *TraesCS6A02G022700* (**Figure 4D**). We further note that this region contains 10,205 bp of gap sequence suggesting this region has been misassembled in IW. This analysis highlights how T4, with its superior genome completeness, resolves genic sequence previously misassembled in IW.

## DISCUSSION

In one critical aspect, the bread wheat genome exemplifies the challenge of eukaryotic genome assembly. Repeats, which remain difficult to assemble, are pervasive in this transposon-rich allopolyploid plant genome. It stands to reason that the accurate and complete resolution of this genome would especially depend on high-quality data and advanced genome assembly techniques. In 2017, we published the first near-complete and contiguous representation of the bread wheat genome (Triticum 3.1), demonstrating that only long DNA sequencing reads are capable of resolving this genome^7^. In our efforts described here, we used Triticum 3.1 as our foundation while leveraging the strengths of the IWGSC reference genome to establish the most complete chromosome-scale and gene-annotated reference assembly yet created for bread wheat.

In scaffolding and annotating our contigs, we created the genomic context needed to quantify and qualify the completeness of the Triticum 4.0 assembly, especially relative to its predecessors. We first observed that, compared to the IWGSC v1.0 assembly, Triticum 4.0 more accurately represents repetitive sequence genome-wide, which also contains more centromeric sequence. Importantly, by successfully resolving repeat sequences, Triticum 4.0 better contextualizes and characterizes protein-coding genes. Because we were able to assemble and anchor a higher proportion of the genome, we were able to place onto chromosomes 2,001 of the 2,691 genes that are unplaced in IW v1.0. Triticum 4.0 also successfully resolves extensive collapsing of gene duplications in IW v1.0, leading to the discovery of 5,799 additional gene copies, and we highlighted a conspicuous example of a newly discovered MADS-box transcription factor gene tandem duplication. Resolving such gene duplications in Triticum 4.0 is consistent with previous studies that estimated gene duplication to be extensive in the bread wheat genome^4^.

## METHODS

### Establishing the initial contig set

We first sought to establish the most complete set of contigs representing the genome of *T. aestivum* Chinese Spring. We started with the Triticum 3.1 contigs (T3)^7^ because they comprise 1 Gbp of additional non-gap sequence compared to the IWGSC v1.0 (IW) reference assembly. However, when establishing a set of contigs for downstream scaffolding, we wanted to ensure that we incorporated any contigs unique to the reference assembly and therefore “missing” from the T3 assembly. To do this, we broke the reference assembly into “contigs” by breaking pseudomolecules at gaps (at least 20 “N” characters). We then aligned these reference contigs (query) to the T3 contigs (reference) using NUCmer (-l 250 -c 500), and filtered them using “delta-filter -1 -l 5000” options to include only reciprocal best alignments at least 5 kbp long^10^. Of reference contigs that were at least 10 kbp in length, if under 25% of that contig was covered by alignments, it was deemed a putative “missing” contig.

We then checked to see if these missing contigs would indeed be covered by alignments produced with more sensitive parameters. The putative missing contigs (query) were aligned again to the T3 assembly with NUCmer, but with a smaller minimum seed and cluster size (-l 50 -c 200). Alignments were filtered as before and if under 25% of a putative missing contig was covered by these more sensitive alignments, they were deemed to be validated as missing from T3. These validated missing IW contigs were combined with the T3 contigs to establish our final set of contigs for downstream scaffolding, which had an N50 length of 230,687 bp and a sum of 15,429,603,425 bp.

### RaGOO scaffolding

We performed two rounds of reference-guided scaffolding with RaGOO. We first used RaGOO to look for false sequence duplications, especially those that could have arisen by incorporating “missing” IW contigs. Though RaGOO usually employs Minimap2^9^ to align query contigs to a reference genome, we used NUCmer in order to produce highly specific alignments. We aligned our contigs (query) to the IW reference genome (reference) using a very large seed and cluster size (-l 500 -c 1000). Such specificity in alignments was necessary to unambiguously order and orient contigs with respect to the highly repetitive allopolyploid reference genome. The resulting delta file was converted to PAF format using Minimap2’s paftools. Next, we ran RaGOO using these alignments rather than the default minimap2 alignments while also specifying a minimum clustering confidence score of 0.4 (-i). We also excluded any unanchored IW sequence from consideration (-e).

To remove false duplication of missing contig sequence, we observed that such duplications would align more than one time in these RaGOO pseudomolecules. Conversely, contigs that were truly “missing” should only align once (perfectly) to their ordered and oriented location in the RaGOO pseudomolecules. We aligned the RaGOO pseudomolecules (query) to the missing IW contigs (reference) with NUCmer (-l 50 -c 200) and filtered alignments with delta-filter (-q -l 5000)^11^. If a missing contig had more than one alignment with coverage at least 50% and percent identity at least 98%, it was deemed to be a false duplicate and removed from the initial contig set. With false duplicates removed, we proceeded with the second round of RaGOO scaffolding which had all of the same specifications as the first round.

Finally, we sought to remove any unanchored contigs that had duplicated sequences amongst the anchored contigs. The same previously described process to remove false duplicates was also used here, except that the RaGOO scaffolds along with unanchored contigs (query) were aligned to the unanchored contigs (reference). Also, the minimum coverage was 75% rather than 50%. Next, scaffolds were polished with POLCA (included in MaSuRCA 3.3.5)^12^. For polishing, we used the Illumina reads from the NCBI SRA accession SRX2994097. POLCA introduced 595,705 bp in substitution corrections and 1,033,593 bp in insertion/deletion corrections. After polishing, the final error rate of the sequence was estimated at less than 0.008% or less than 1 error per 10,000 bases. T4/IW dotplots were made by aligning the polished T4 assembly (query) to the IW reference assembly (reference) with NUCmer (-l 500 -c 1000). To produce a single dotplot depicting all 21 chromosomes, alignments were filtered with delta-filter (-l 50000 -1). These alignments were then provided to mummerplot (--fat --layout) to produce the final dotplots. The same process was used to create dotplots for individual chromosomes, though alignments less than 10 kbp, rather than 50 kbp, were removed with delta-filter.

### Shared k-mer frequency distribution

101-mers were counted in T4 and IW using KMC (v3.1.0, -ci1 -cx10000 -cs10000)^18^. 101-mers shared by T4 and IW were then extracted with kmc_tools “simple” using the intersection function. The 101-mer copy frequency distribution of these shared *k*-mers in both T4 and IW (-ocleft and - ocright) was then plotted in **Figure 3**.

### Centromere annotation

We annotated centromere sequence in both T4 and IW using a similar approach as that described in the original IW publication^3^. First, publicly available Chinese Spring CENH3 ChIP-seq data (SRR1686799) was downloaded from the European Nucleotide Archive^19^. Reads were then trimmed with cutadapt (v1.18, -a AGATCGGAAGAG) and aligned to T4 and IW with bwa mem (v0.7.17-r1198-dirty)^20,21^. Alignments with a mapq score less than 20 were removed and the remaining alignments were compressed and sorted with samtools view and samtools sort respectively^22^. Alignments were then counted in 100 kbp non-overlapping windows along the T4 and IW genomes using bedtools makewindows and bedtools coverage (v2.29.2)^23^. Any group of two or more consecutive windows with greater than or equal to three times the genomic average coverage was considered putative centromere sequence, and any such intervals within 500 kbp were merged together. The final list of all annotated centromeres for both T4 and IW is found in **Table S1**. Whenever we chose a single centromeric interval for each chromosome (e.g. as depicted in **Figure 2**), the largest interval for each chromosome was chosen. In **Figure 2**, we use centromere coordinates as they are defined in the original IW publication to build our IW ideogram. However, when we report in our results that T4 has 38.8 Mbp more centromeric sequence than IW, we use the IW centromere annotations established in this paper to draw that comparison. Our IW centromere coordinates match closely with those previously defined, with our annotations only containing 3.9 Mbp more sequence.

### Chloroplast and mitochondria genome assembly

We took the first 20 million Illumina read pairs from the SRR5815659 accession and assembled them with megahit (v1.2.8)^24^. The resulting assembly contained 145,887 contigs (74.41Mb) with lengths ranging between 200bp and 56,565 bp. Then we aligned these contigs to the *Triticum aestivum* reference chloroplast sequence (NC_002762.1) using NUCmer (with --maxmatch switch to align to repeats) and filtered the alignments with delta-filter, keeping the best hits to the reference NC_002762.1. The reference was covered completely by alignments of only five contigs. Then, we aligned these contigs to each other with NUCmer (--maxmatch --nosimplify) and used the alignments to manually order and orient them into a single chloroplast sequence scaffold.

To establish a mitochondria sequence, we aligned the megahit contigs discussed above to the *Triticum aestivum* mitochondria reference sequence (MH051716) with NUCmer (--maxmatch). We then filtered the alignments with delta-filter, keeping the best matches to the MH051716 reference. This revealed 43 non-chloroplast contigs of least 500 bp in length that matched best to the mitochondria reference. We then ordered and oriented these 43 contigs using RaGOO (v1.1), setting the minimum alignment length to 500 bp. The chloroplast and mitochondria sequence are included in our data submission to NCBI.

### Genome annotation

We used the gffutils v0.10.1 python library to read the reference annotation and sequence and extract complete gene sequences (introns and exons). We then built a separate BLAST nucleotide database of each T4 chromosome^14^. Genes were aligned to their same chromosome in T4 using BLASTN v.2.9.0 (-soft_masking False -dust no -word_size 50 -gap_open 3 -gapextend 1 - culling_limit 10). The blast hits were filtered to include only those that contained one or more exons. For each gene, the optimal exon alignments were chosen according to sequence identity and concordance with the exon/intron structure of the gene model in IW. These alignments were used to define the boundaries of each exon, transcript, and gene in T4. We excluded any transcripts that did not map with at least 50% alignment coverage. Any genes without at least 1 mapped isoform were then aligned against the entire T4 genome using BLASTN with the same parameters and placed given they did not overlap an already placed gene.

To place the chrUn genes, we aligned the genes to the entire T4 genome using the same parameters. We excluded any transcripts that did not meet the 50% alignment coverage threshold or overlapped an already annotated gene.

To find additional gene copies we aligned all genes to the complete T4 genome using BLAST v2.9.0 (-soft_masking False -dust no -word_size 50 -gap_open 3 -gapextend 1 -culling_limit 100, qcov_hsp_perc 100). The notable differences in these parameters are qcov_hsp_perc which requires 100% query coverage, and culling_limit which has been increased from 10 to 100 to increase the number of reported alignments for genes with a highly increased copy number. We excluded any alignments that did not have 100% exonic sequence identity or overlapped a previously placed gene. We used gffread to filter out genes with non-canonical splice sites (https://github.com/gpertea/gffread).

## ACKNOWLEDGMENTS

This work was supported in part by NIH under grants R01-HG006677 and R35-GM130151, and by the USDA National Institute of Food and Agriculture under grant 2018-67015-28199.

## DATA AVAILABILITY

The Triticum 4.0 assembly and annotations are available at www.ncbi.nlm.nih.gov/bioproject/PRJNA392179(GenBank accession: GCA_002220415.3)

## AUTHOR CONTRIBUTIONS

S.S. designed the project. M.A., A.S., D.P., A.Z., and S.S. designed analysis. M.A., A.S., D.P., A.Z., and S.S. analyzed data. M.A., A.S., and S.S. wrote the manuscript. All authors read and approved the final manuscript.

---

**Figure S1.**
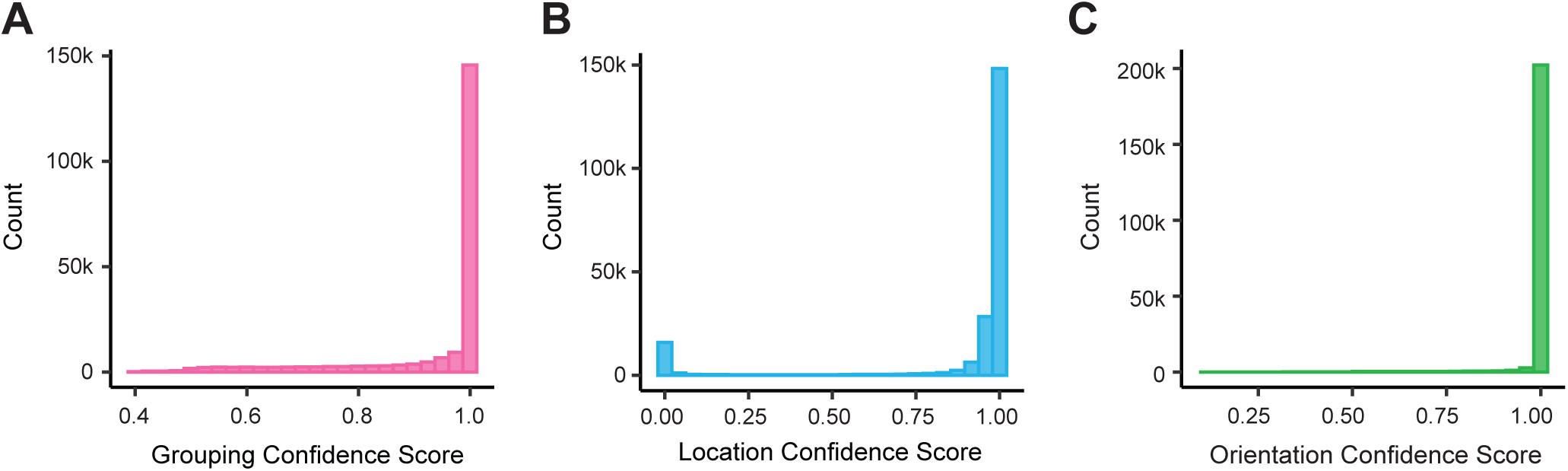
RaGOO confidence score distributions. Histograms depicting the distributions for “grouping”, “location”, and “orientation” confidence scores. Every input contig is assigned a “confidence score” for the 3 scaffolding steps. Scores are between 0 and 1, with higher scores corresponding to less ambiguous contig ordering (“grouping” and “location”) and orientation (“orientation”).

**Figure S2.**
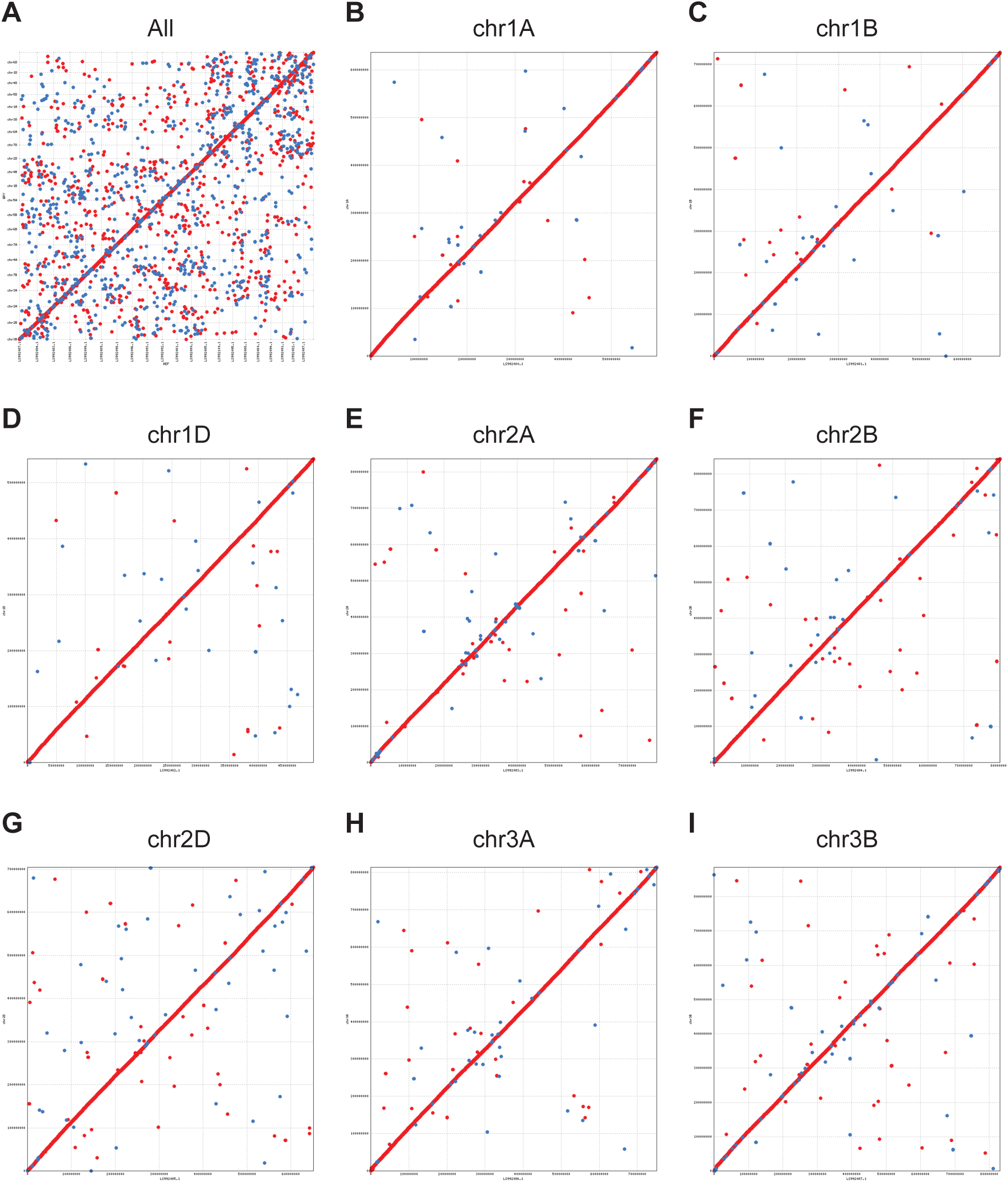
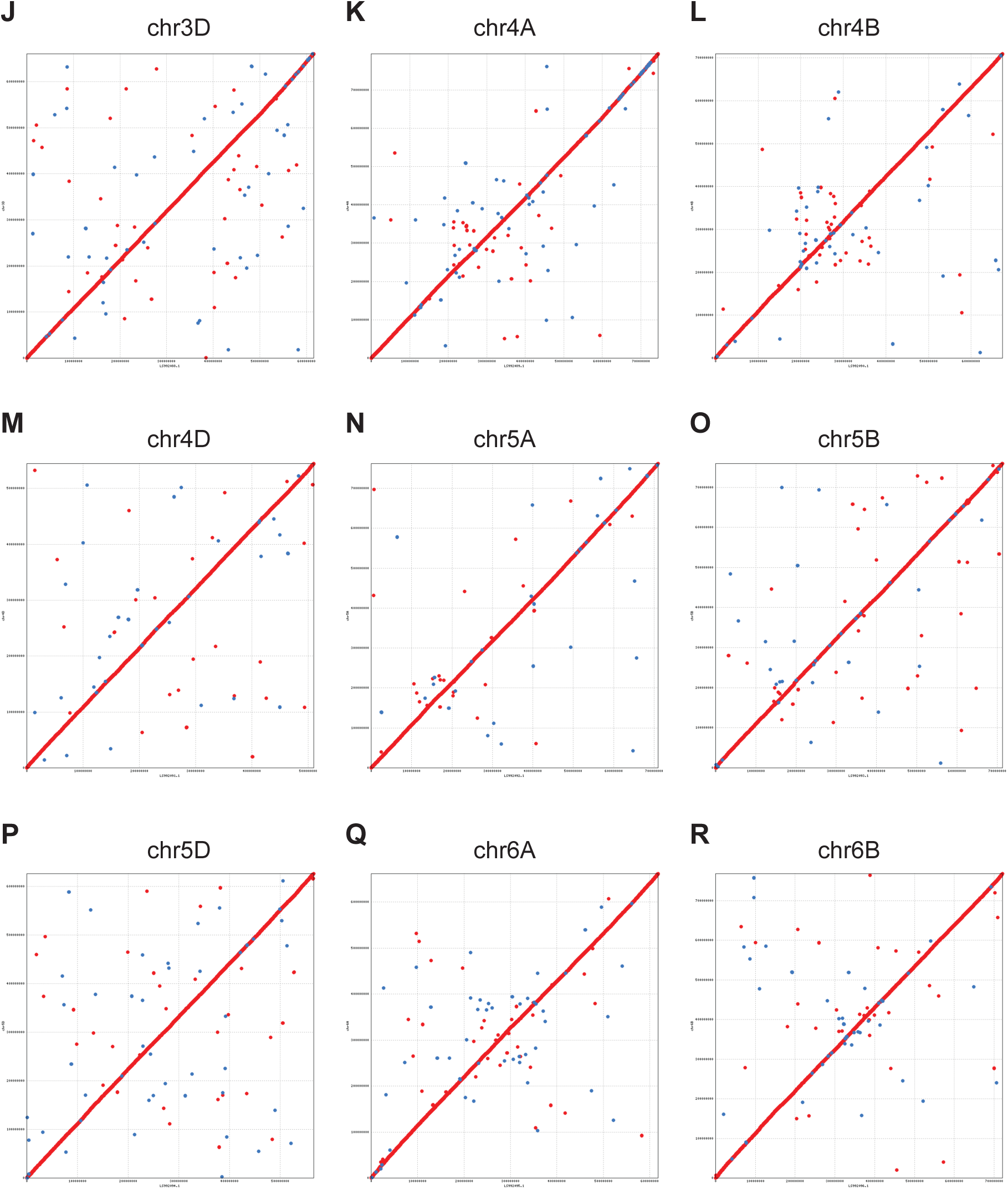
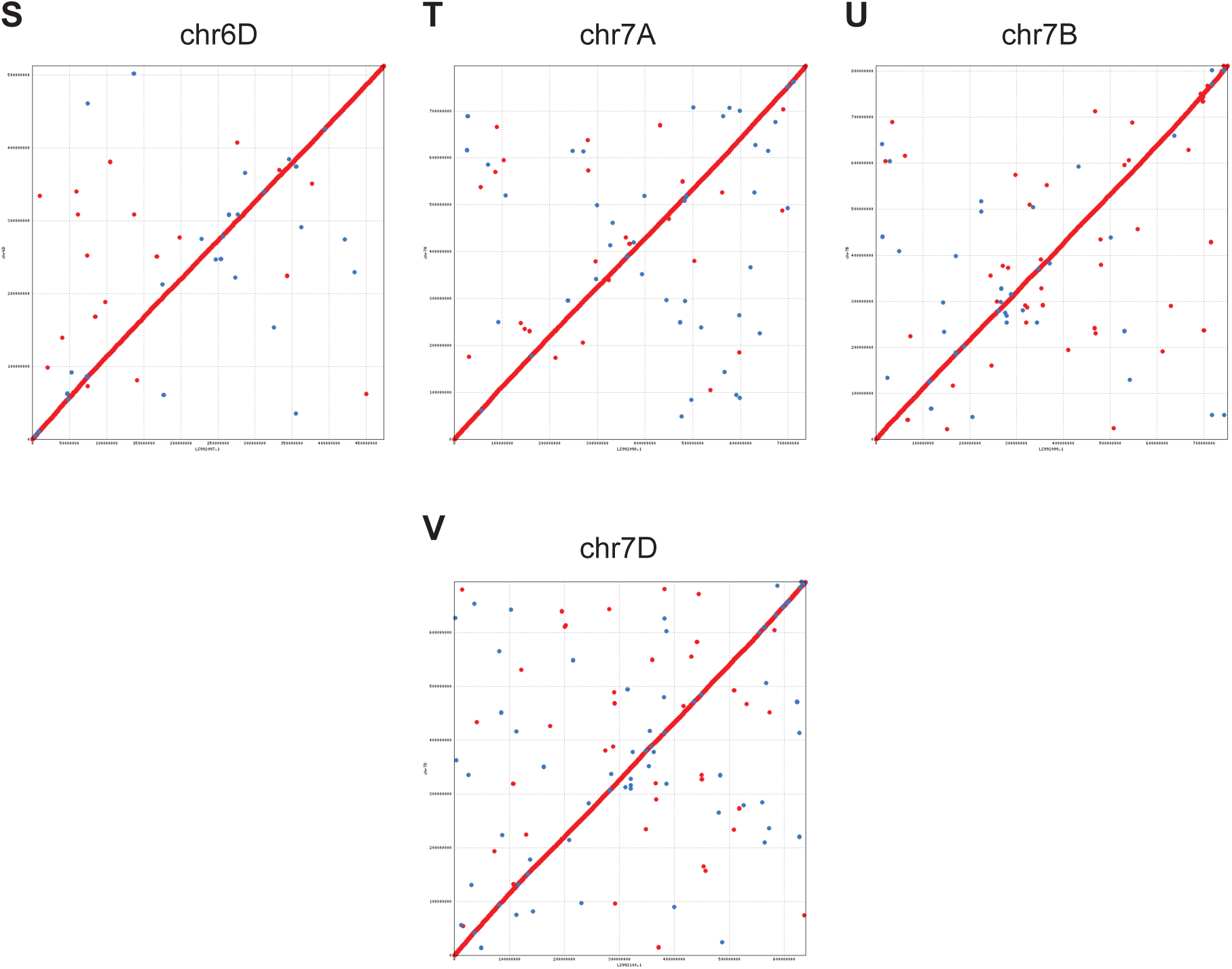
Triticum 4.0 genome assembly dotplots. Triticum 4.0 (QRY) dotplots with respect to the IWGSC v1.0 reference assembly (REF). Panel **(A)** shows the genome-wide dotplot, while panels **(B-V)** show dotplots for individual chromosomes. The order of panels (B-V) corresponds to the lexicographically sorted order of the chromosome names (e.g. chr1A). As expected, no large-scale structural differences are observed between the two assemblies.

**Figure S3.**
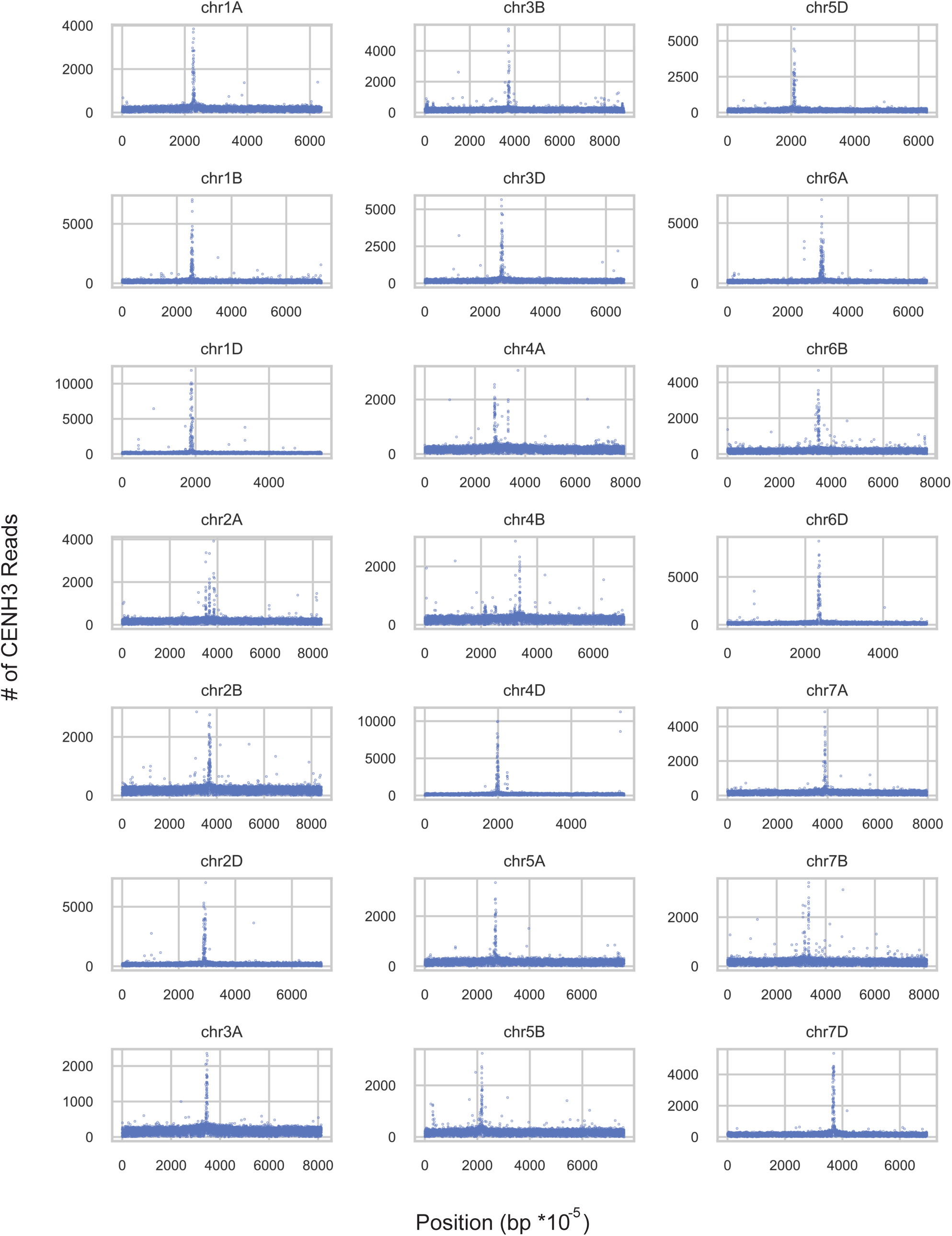
Triticum 4.0 CENH3 ChIP-seq coverage. Scatter plots depicting the number of CENH3 ChIP-seq read alignments (y-axis) in 100 kbp non-overlapping windows (x-axis) of the Triticum 4.0 genome assembly. Peaks in these scatterplots correspond to putative centromere sequences.

**Figure S4.**
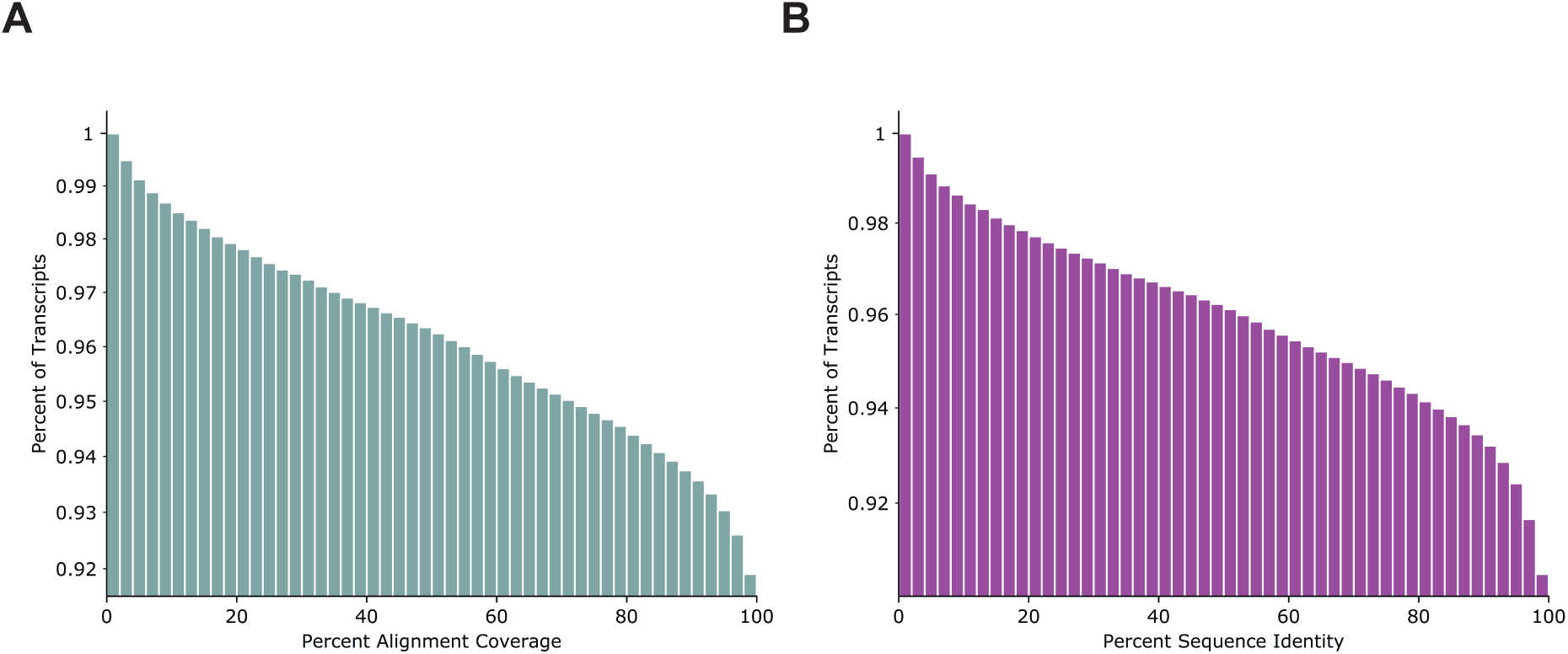
Alignment coverage and sequence identity cumulative distribution. **(A)** Cumulative distribution showing how much of the IW transcripts map onto T4. The Y axis shows the fraction of transcripts with percent coverage greater than or equal to coverage on the X axis. **(B)** Cumulative distribution showing the sequence identity of IW transcripts mapped onto T4. The Y axis shows the fraction of transcripts with sequence identity greater than or equal to sequence on the X-axis.

## REFERENCES

1. Dubcovsky, J. & Dvorak, J. Genome plasticity a key factor in the success of polyploid wheat under domestication. Science vol. 316 1862–1866 (2007).

2. Arumuganathan, K. & Earle, E. D. Nuclear DNA content of some important plant species. Plant Mol. Biol. Report. 9, 208–218 (1991).

3. Appels, R. et al. Shifting the limits in wheat research and breeding using a fully annotated reference genome. Science (80-.). 361, (2018).

4. Mayer, K. F. X. et al. A chromosome-based draft sequence of the hexaploid bread wheat (Triticum aestivum) genome. Science (80-.). 345, 1251788–1251788 (2014).

5. Clavijo, B. J. et al. An improved assembly and annotation of the allohexaploid wheat genome identifies complete families of agronomic genes and provides genomic evidence for chromosomal translocations. Genome Res. 27, 885–896 (2017).

6. Chapman, J. A. et al. A whole-genome shotgun approach for assembling and anchoring the hexaploid bread wheat genome. Genome Biol. 16, (2015).

7. Zimin, A. V. et al. The first near-complete assembly of the hexaploid bread wheat genome, Triticum aestivum. Gigascience (2017) doi:10.1093/gigascience/gix097.

8. Alonge, M. et al. RaGOO: Fast and accurate reference-guided scaffolding of draft genomes. Genome Biol. 20, (2019).

9. Li, H. Minimap2: pairwise alignment for nucleotide sequences. Bioinformatics 34, 3094–3100 (2018).

10. Kurtz, S. et al. Versatile and open software for comparing large genomes. Genome Biol. 5, R12 (2004).

11. Marçais, G. et al. MUMmer4: A fast and versatile genome alignment system. PLoS Comput. Biol. (2018) doi:10.1371/journal.pcbi.1005944.

12. Zimin, A. V. & Salzberg, S. L. The genome polishing tool POLCA makes fast and accurate corrections in genome assemblies. bioRxiv 2019.12.17.864991 (2019) doi:10.1101/2019.12.17.864991.

13. Schatz, M. C., Delcher, A. L. & Salzberg, S. L. Assembly of large genomes using second-generation sequencing. Genome Research (2010) doi:10.1101/gr.101360.109.

14. Altschul, S. F., Gish, W., Miller, W., Myers, E. W. & Lipman, D. J. Basic local alignment search tool. J. Mol. Biol. (1990) doi:10.1016/S0022-2836(05)80360-2.

15. Coen, E. S. & Meyerowitz, E. M. The war of the whorls: genetic interactions controlling flower development. nature.comPaperpile https://www.nature.com/articles/353031a0(1991).

16. Ng, M., Genetics, M. Y.-N. R. & 2001, undefined. Function and evolution of the plant MADS-box gene family. nature.comPaperpile.

17. Soyk, S. et al. Duplication of a domestication locus neutralized a cryptic variant that caused a breeding barrier in tomato. Nature Plants vol. 5 471–479 (2019).

18. Kokot, M., Dlugosz, M. & Deorowicz, S. KMC 3: counting and manipulating k-mer statistics. Bioinformatics (2017) doi:10.1093/bioinformatics/btx304.

19. Guo, X. et al. De Novo Centromere Formation and Centromeric Sequence Expansion in Wheat and its Wide Hybrids. PLoS Genet. (2016) doi:10.1371/journal.pgen.1005997.

20. Martin, M. Cutadapt removes adapter sequences from high-throughput sequencing reads. EMBnet.journal (2011) doi:10.14806/ej.17.1.200.

21. Li, H. & Durbin, R. Fast and accurate short read alignment with Burrows-Wheeler transform. Bioinformatics 25, 1754–1760 (2009).

22. Li, H. et al. The Sequence Alignment/Map format and SAMtools. Bioinformatics 25, 2078–2079 (2009).

23. Quinlan, A. R. & Hall, I. M. BEDTools: a flexible suite of utilities for comparing genomic features. Bioinformatics 26, 841–842 (2010).

24. Li, D., Liu, C. M., Luo, R., Sadakane, K. & Lam, T. W. MEGAHIT: An ultra-fast single-node solution for large and complex metagenomics assembly via succinct de Bruijn graph. Bioinformatics (2015) doi:10.1093/bioinformatics/btv033.

